# Anterior fontanelle size among term neonates on the first day of life born at University of Gondar Hospital, Northwest Ethiopia

**DOI:** 10.1101/385906

**Authors:** Mohammed Oumer, Edengenet Guday, Alemayehu Teklu, Abebe Muche

## Abstract

**Background:** Anterior fontanelle is the largest, prominent and most important fontanelle, which is used for clinical evaluation. It is mainly characterized by its size and shape variation and is possibly influenced by gender, race and genetics. Understanding the variation of anterior fontanelle is used for recognition of different medical disorders and abnormal skeletal morphogenesis.

**Objective:** To determine the mean size of anterior fontanelle among term neonates on the first day of life born at University of Gondar Hospital, Gondar town, Northwest Ethiopia, 2018

**Methods:** Descriptive cross sectional study design was undertaken in 384 term and apparently healthy neonates, using standard methods. Descriptive analysis, student t-test, one way ANOVA and Pearson correlation coefficient were implemented.

**Results:** In this study, the mean size of anterior fontanelle in term neonates was 3.00 ± 0.62 cm (range 1.70 – 5.50 cm). The mean size of anterior fontanelle was 3.10 ± 0.66 cm for males, and 2.88 ± 0.57 cm for females. There was statistically significant difference in anterior fontanelle size in neonates of different genders (*p*<0.001), mode of delivery (*p*<0.001) and duration of labour (*p*=0.006). However, the size of anterior fontanelle was not significantly affected by the birth order, onset of labour and sociodemographic variables of the mother except occupation of the mother (p=0.01). There was a significant positive correlation between the mean size of anterior fontanelle with birth weight (*r*=0.11; *p*=0.04) and head circumference (*r*=0.17; *p*=0.001).

**Conclusions:** At term, male neonates had significantly larger anterior fontanelle than female neonates and anterior fontanelle size has a direct relationship with birth weight and head circumference.

## Introduction

Fontanelles are fibrous gaps occurring when more than two cranial bones are juxtaposed or it can be defined as a place where two or more sutures meet (1–4). The word fontanelle is derived from the Latin word, Fonticulus and the old French word, Fontaine, meaning a little fountain or spring (2–5). In the newborn skull six fontanelles can be identified, namely anterior, posterior, two mastoid and two sphenoid fontanelles (6–8). Out of six, the most prominent and the largest fontanelle is rhomboid anterior fontanelle (AF) situated between the two frontal and two parietal bones (8). AF allows growth of the brain in relative to skull bone growth (2, 4, 9). The bones of the skull overlap each other during labour time for successful delivery; however, the molding process of the skull will be resolved after three to five days of birth (2, 4). The average time of AF closure is 18 months, but often closes by 12 months (8, 10, 11). The variation in size, shape and closure time is a key feature of normal AF (5, 8). Until recently, AF measurement has not been routinely performed as part of the newborn examination, even if, the clinical value of AF examination is crucial (6).

AF size has been utilized as evidence of altered intracranial pressure, an index of the rate of development and ossification of the calvarium (5). It is also an indicator to various medical disorders and abnormal skeletal morphogenesis (2, 4–6, 12).

The diagnosis of an abnormal fontanelle e.g. bulging, sunken fontanelle, large size, small size, early closure and delayed closure requires an understanding of the wide variation of normal fontanelle. Knowledge of AF size is crucial to determine many disorders. Delayed closure or large size of the AF can be associated with multiple diseases. Of them, skeletal disorders, chromosomal defects and dysmorphogenesis syndromes, endocrine disorders, drug and toxin exposure, fetal hydantoin syndrome, aminopterin induced malformations, congenital infections (e.g. rubella and syphilis) and aluminum toxicity (4, 8, 10).

Increased intracranial pressure is the most common cause of bulging, enlarged and delayed closure of the AF. A sunken AF is the sign of dehydration (4, 8). In cardiac diseases (a full fontanel, which is tense and visibly elevated may support the diagnosis of a heart failure), while in intrauterine growth retardation (neonates with small for date of gestational age) may show the largest AF (3, 13).

Physical examination of AF size along with well child care in neonates and infants is highly recommended medical practices in Neonatology and Pediatrics. It gives useful information to follow the developmental status of the child as well as general state of health and can be considered as an index of cranial growth and development during the prenatal and postnatal period. Any developmental alteration in AF growth presumably is an indicator of an abnormal growth (2, 3, 6, 14).

In Ethiopia, like any other developing countries, published literature regarding size of AF is scarce. In order to establish a national age related standard for AF size, further study in different parts of Ethiopia was recommended in the study done in Addis Ababa on infant's age between 3 days and 270 days in 2004 (3). Hence, the present study is undertaken in attempt to fill the gap by providing information on the size of AF of neonates living in Gondar.

## Methods and materials

### Study area and period

An institution based descriptive cross sectional study design was undertaken and the study was conducted at Maternity and Neonatal Ward, University of Gondar Hospital (UoGH), Gondar town from October 01, 2017 to February 15, 2018.

### Sample size and sampling technique

The sample size was calculated using a single population proportion formula; by considering p-value 50 %, at 95% confidence interval (CI), 5% margin of error and 5% non-response rate.

Mothers of the neonates were selected randomly at the time period of 6:00 am-2:00 pm. All apparently healthy term neonates on the first day of life were included in the study. Neonates with gestational age between 37-42 weeks (appropriately grown neonates), birth weight between 2500 gm and 4000 gm, and neonates free from any medical and physical disorders, that affects AF size, were included in this study.

A total of 384 term neonates (206 males and 178 females), born at the time of data collection were included in the study.

### Study variables

AF Size is a dependent variable

Independent variables include socio demographic variables; maternal age, place of residence, marital status, educational status, occupation and monthly income of the mother. And, pregnancy, labour, outcome and newborn related variables; birth order, onset of labour, duration of labour, mode of delivery, gender of neonate, birth weight and head circumference.

### Data collection tools, techniques and procedures

Data were collected using a checklist with direct face to face interview with the mothers. The main questions were maternal socio demographic variables, and pregnancy, labour, outcome and newborn related variables. The data collection checklist was first prepared in English and translated to Amharic and then back to English to check for consistency of meaning.

The conditions during delivery, maternal socio demographic variables, birth order, onset of labour, duration of labour and mode of delivery (Spontanious vertex delivery or Ceserian section) of the neonate were obtained from the mother and attending physician. Birth weight, head circumference, and anterior posterior and transverse dimension of AF were measured using non-elastic (non-stretchable) plastic tape within 24 hours of birth. All measurements were in centimeters and grams. The AF size was measured by using Popich and Smith method (2, 3, 6, 8). The Popich and Smith method is the simplest, practical and acceptable in clinical settings (Fig. 1). The extent of the AF was determined by inserting the index finger in turn into each of the four corners (vertices) (Fig. 2) and a small circular dot was marked with washable ink on the skin immediately distal to the finger. Then, a piece of white paper was firmly pressed over the fontanel so that the four dots were transferred onto the paper as traced and marked according to Davies et al. in 1975 (15). The distance between the anterior and posterior points and between the transversal points were measured and recorded with a fresh ruler, with an accuracy of closest millimeter. The average (mean) of anterior posterior dimension (length) and transverse dimension (width) was considered as the size of AF (12). The fontanelle was examined while the neonate is calm and held in both supine and upright position (4, 8).

**Figure 1.**
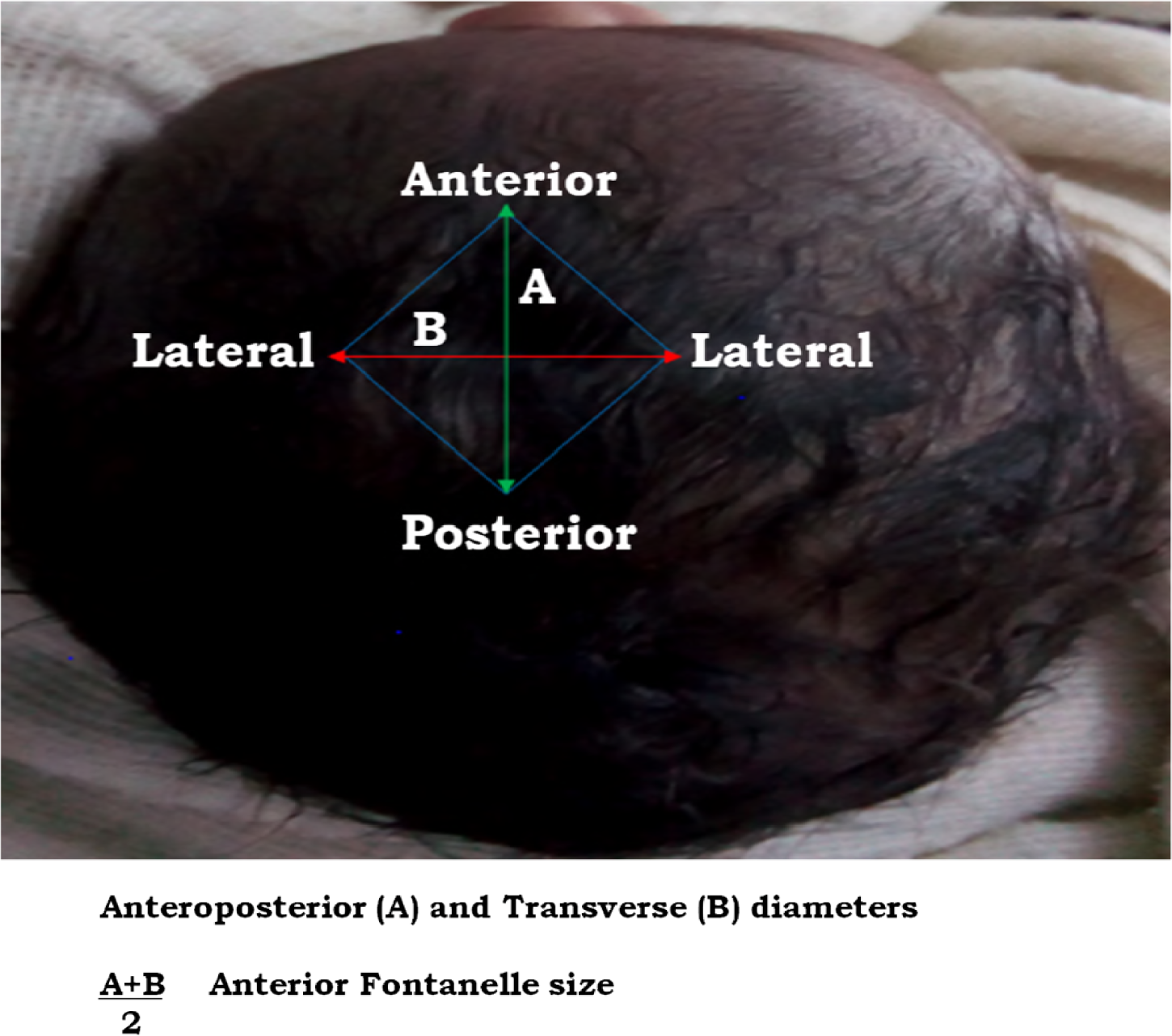
Method of measurements of mean size of AF. 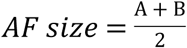 (based on Popich and Smith).

**Figure 2.**
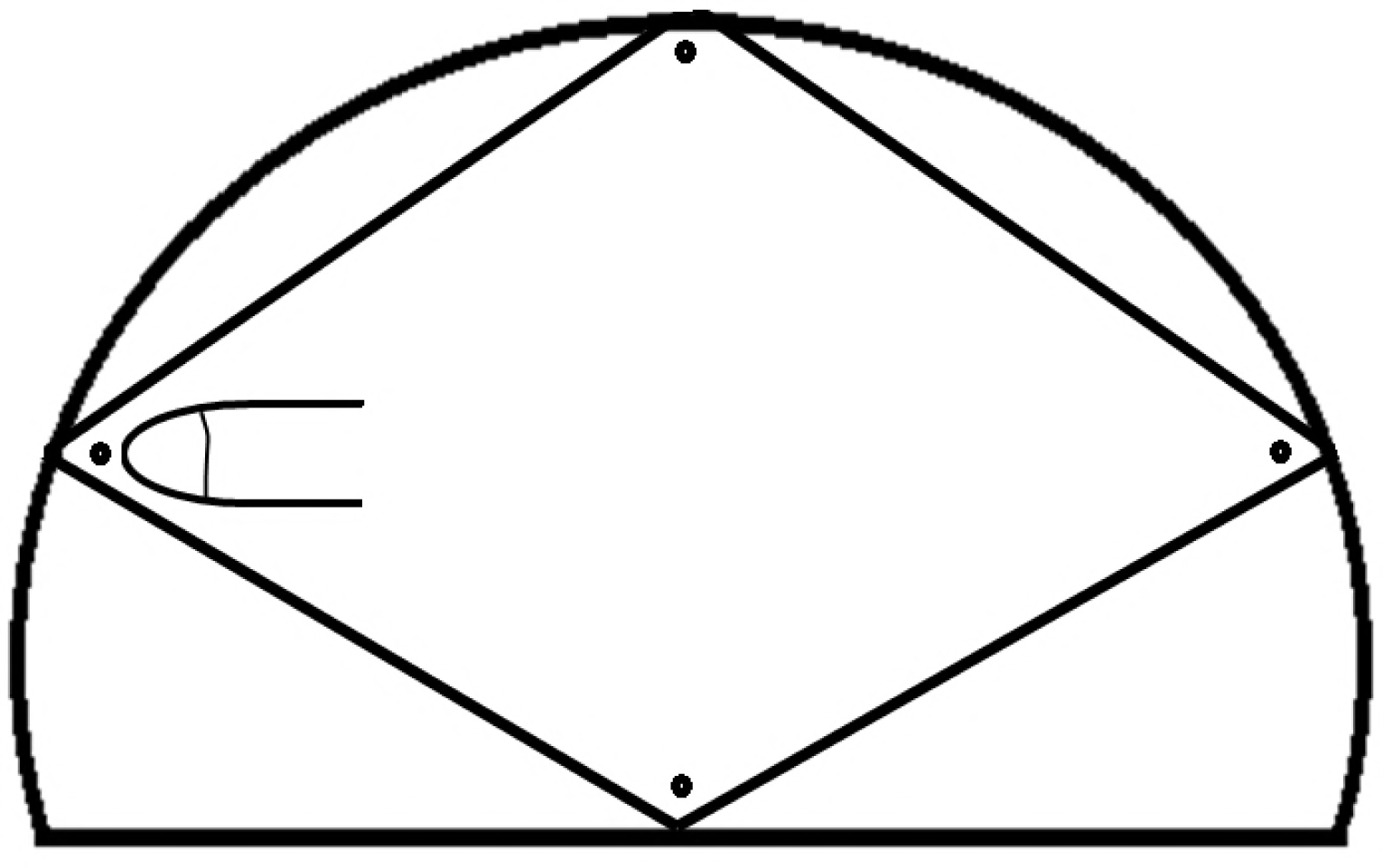
Measurement of AF size in the neonate (determining four vertices).

**Figure 3.**
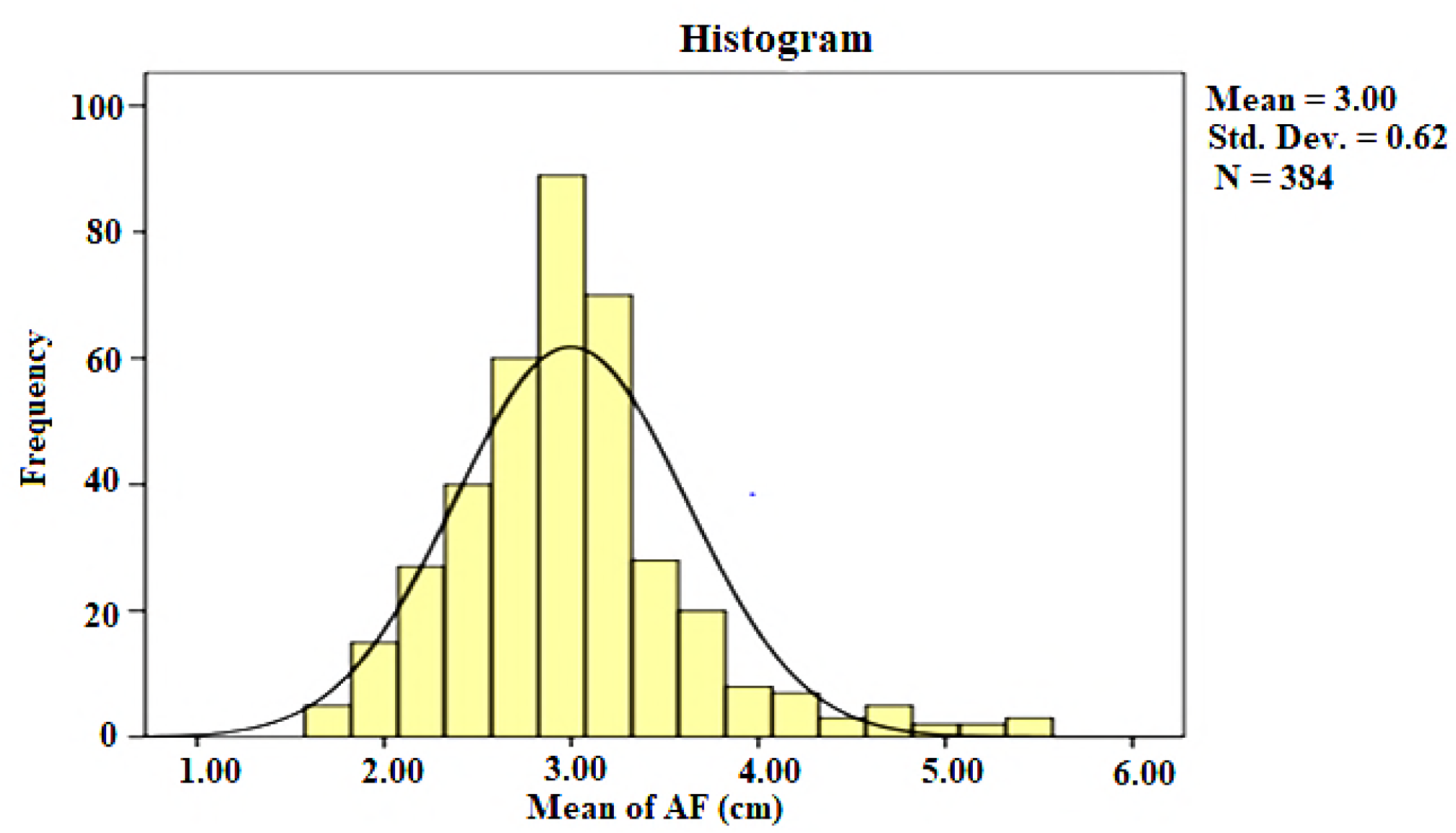
Distribution of AF size among term neonates of UoGH, Gondar town, Northwest Ethiopia, 2018

For each neonate, head circumference was measured with calibrated non elastic (non stretchable) plastic tape. The tape was placed around the neonate’s head at the same level on each side crossing the forehead superior to the supra orbital ridges and passing over the occipital prominence posteriorly with an accuracy of closest millimeter. In order to measure the weight of neonate, a balance beam neonate scale was used with the accuracy of closest gram.

All infection prevention precautions standards were used during the time of measurement. Standard precautions were also applied for measuring equipments.

Materials like fresh ruler, washable ink, a balance beam neonate scale and calibrated non elastic plastic tape were used to measure parameters. All measurements were recorded on the checklist designed for this study.

### Data processing and analysis

The collected data were checked for completeness, accuracy and clarity. The collected data were entered, cleaned, validated and analyzed by using SPSS (Statistical Package for Social Science) version 20. Descriptive analysis was done to describe the frequency and percentage of dependent and independent variables. The mean and standard deviation (S.D.) of the AF size were analyzed for the two genders. Two-tailed student t-test (independent) was used to compare the difference in AF size between genders, mode of delivery, onset of labour and place of residence. One way ANOVA (Analysis of Variance) test was utilized to compare neonates among groups of birth order, duration of labour, maternal age, marital status, educational status, occupation and monthly income of the mother in the mean difference in their AF size. The Pearson correlation coefficient was used to identify the correlation between AF size with birth weight and head circumference. Percentiles of the AF size were also drawn. A P-value less than 0.05 were considered to be statistically significant.

### Ethical consideration

Ethical clearance was obtained from the School of Medicine, College of Medicine and Health Sciences, University of Gondar Ethical review committee. The purpose of the study was described to the mothers of all neonates, and a written or oral informed consent was taken from the neonate’s mother or father. In addition, all information obtained from them was secured and kept confidential. To ensure confidentiality, the names were avoided from the checklist. All data involving measurement were collected without any risk or harm to the neonates.

## Results

### Socio-demographic characteristics of mothers

A total of 384 mothers of neonates were involved in the study (response rate = 100%), of whom 284 (74.0%) were urban residents. The majority of the women were married 366 (95.3%) and in the age group of 26 to 30 years old 166 (43.2%). About half of respondents were with educational status of 6 to 12 grades 193 (50.3%) and housewives in occupation 201 (52.3%) (Table 1).

**Table 1.**
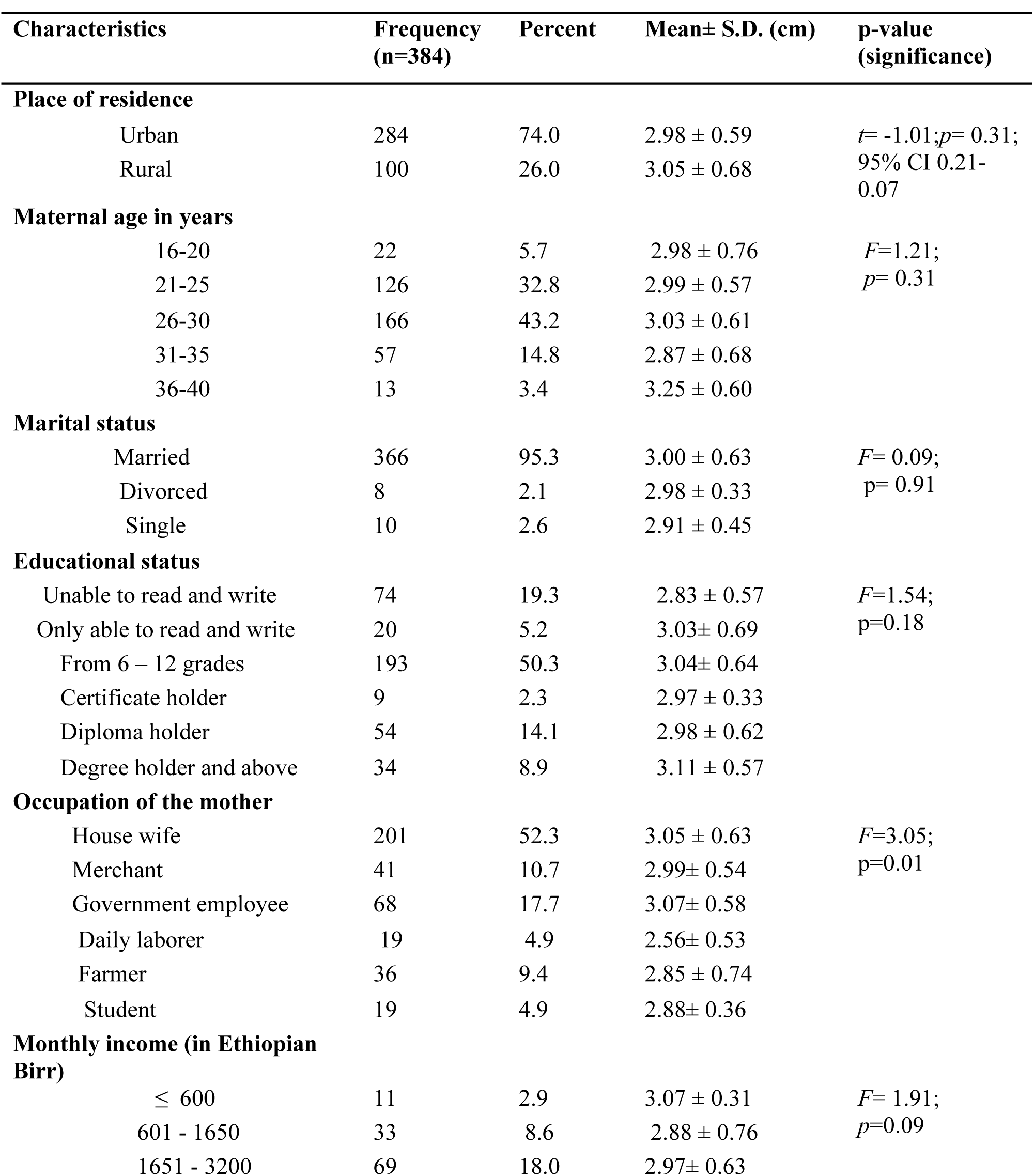
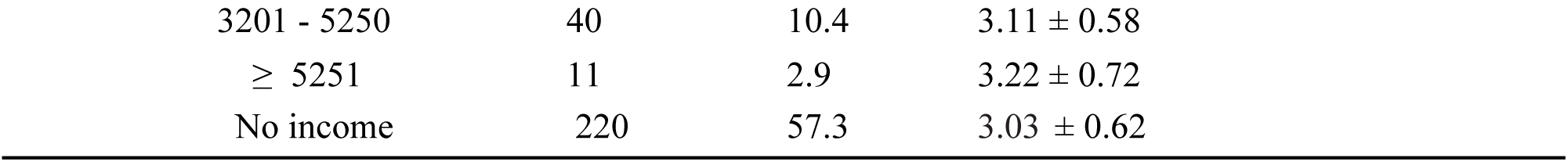
Socio-demographic characteristics of mothers and its mean, S.D. and p-value of UoGH, Gondar town, Northwest Ethiopia, 2018

| Characteristics | Frequency<br>(n=384) | Percent | Mean± S.D. (cm) | p-value<br>(significance) |
| --- | --- | --- | --- | --- |
| Place of residence |  |  |  |  |
| Urban | 284 | 74.0 | 2.98 ± 0.59 | *t*= -1.01;*p*= 0.31;<br>95% CI 0.21-0.07 |
| Rural | 100 | 26.0 | 3.05 ± 0.68 |  |
| Maternal age in years |  |  |  |  |
| 16-20 | 22 | 5.7 | 2.98 ± 0.76 | *F*=1.21;<br>*p*= 0.31 |
| 21-25 | 126 | 32.8 | 2.99 ± 0.57 |  |
| 26-30 | 166 | 43.2 | 3.03 ± 0.61 |  |
| 31-35 | 57 | 14.8 | 2.87 ± 0.68 |  |
| 36-40 | 13 | 3.4 | 3.25 ± 0.60 |  |
| Marital status |  |  |  |  |
| Married | 366 | 95.3 | 3.00 ± 0.63 | *F*= 0.09;<br>*p*= 0.91 |
| Divorced | 8 | 2.1 | 2.98 ± 0.33 |  |
| Single | 10 | 2.6 | 2.91 ± 0.45 |  |
| Educational status |  |  |  |  |
| Unable to read and write | 74 | 19.3 | 2.83 ± 0.57 | *F*=1.54;<br>*p*=0.18 |
| Only able to read and write | 20 | 5.2 | 3.03± 0.69 |  |
| From 6 – 12 grades | 193 | 50.3 | 3.04± 0.64 |  |
| Certificate holder | 9 | 2.3 | 2.97 ± 0.33 |  |
| Diploma holder | 54 | 14.1 | 2.98 ± 0.62 |  |
| Degree holder and above | 34 | 8.9 | 3.11 ± 0.57 |  |
| Occupation of the mother |  |  |  |  |
| House wife | 201 | 52.3 | 3.05 ± 0.63 | *F*=3.05;<br>*p*=0.01 |
| Merchant | 41 | 10.7 | 2.99± 0.54 |  |
| Government employee | 68 | 17.7 | 3.07± 0.58 |  |
| Daily laborer | 19 | 4.9 | 2.56± 0.53 |  |
| Farmer | 36 | 9.4 | 2.85 ± 0.74 |  |
| Student | 19 | 4.9 | 2.88± 0.36 |  |
| Monthly income (in Ethiopian Birr) |  |  |  |  |
| ≤ 600 | 11 | 2.9 | 3.07 ± 0.31 | *F*= 1.91;<br>*p*=0.09 |
| 601 - 1650 | 33 | 8.6 | 2.88 ± 0.76 |  |
| 1651 - 3200 | 69 | 18.0 | 2.97± 0.63 |  |

|  |  |  |  |
| --- | --- | --- | --- |
| 3201 - 5250 | 40 | 10.4 | $3.11 \pm 0.58$ |
| $\geq 5251$ | 11 | 2.9 | $3.22 \pm 0.72$ |
| No income | 220 | 57.3 | $3.03 \pm 0.62$ |

### Anterior Fontanel size

The mean size of AF for the study population was 3.00 cm (± Standard deviation (S.D.) 0.62 cm with a range of 1.70 – 5.50 cm (Fig. 4).

46.9% and 44.3% of neonate had AF size between 2.0 and 2.99 cm, and 3.0 – 3.99 cm, respectively. Of the total study subjects 11 (2.9%) had AF size of less than 2.0 cm and 6 (1.6%) of them had AF size of greater or equal to 5.0 cm (Table 2).

**Table 2.**
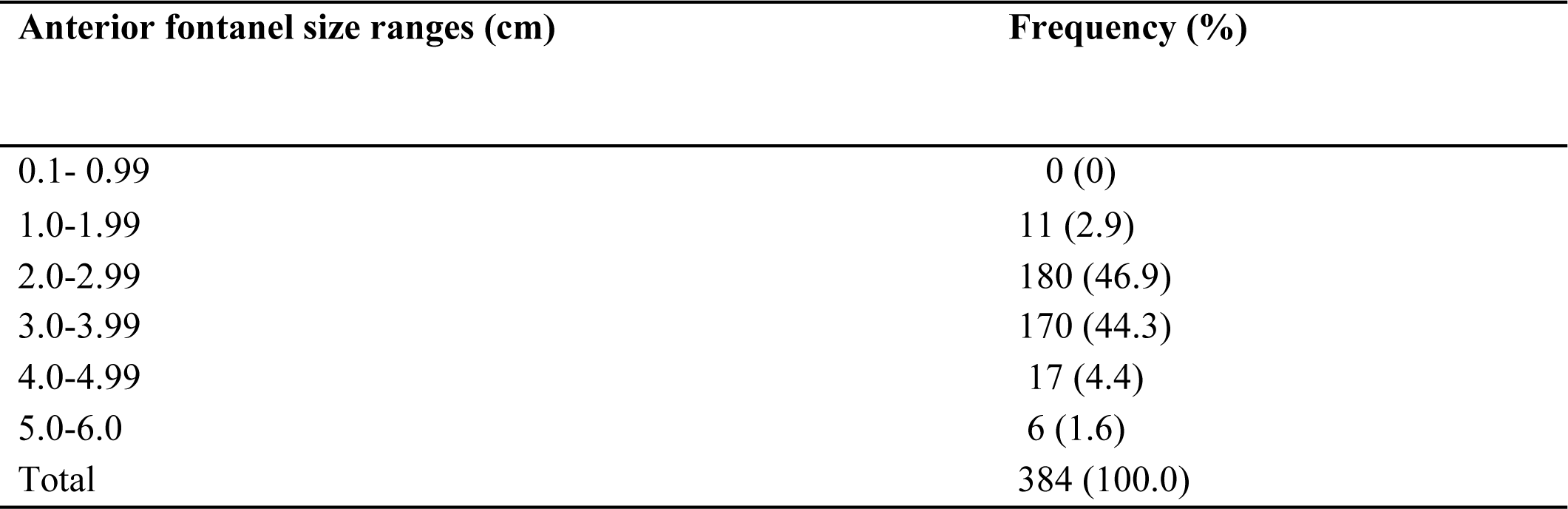
The frequency distribution of AF size among 384 term neonates of UoGH, Gondar town, Northwest Ethiopia, 2018

Based on the percentile distribution, less than five percent of neonates had an AF size below 2.05 cm and five percent had AF size above 4.24 cm. This indicated 90% of the sample had AF size between 2.05 and 4.24 cm (Table 3).

**Table 3.** Percentiles of AF size (cm) among 384 term neonates of UoGH, Gondar town, Northwest Ethiopia, 2018

| Percentiles |  |  |  |  |  |  |  |
| --- | --- | --- | --- | --- | --- | --- | --- |
| At | 5<sup>th</sup> | 10<sup>th</sup> | 25<sup>th</sup> | 50<sup>th</sup> | 75<sup>th</sup> | 90<sup>th</sup> | 95<sup>th</sup> |
| birth | 2.05 | 2.30 | 2.65 | 3.00 | 3.25 | 3.70 | 4.24 |

Of 384 neonates, 206 (53.6%) were males. The mean size of AF was calculated in both genders: 3.10 ± 0.66 cm for males, and 2.88 ± 0.57 cm for females. A statistically significant difference in mean AF size between males and females were found (*t*=-3.69; *p*<0.001).

The mean AF sizes of neonates among groups of birth order and onset of labour were compared for mean difference; nonetheless, statistical difference among these variables were not detected (*p*>0.05) (Table 4).

**Table 4.** Distribution, mean, S.D. and p-value of pregnancy and labour related characteristics among term neonates of UoGH, Gondar town, Northwest Ethiopia, 2018

| Characteristics | Frequency<br>(n=384) | Percent<br>(%) | Mean± S.D. (cm) | p- value<br>(significance) |
| --- | --- | --- | --- | --- |
| Birth order |  |  |  |  |
| 1-2 | 314 | 81.8 | 2.97 ± 0.62 | F= 1.25;<br>p= 0.29 |
| 3-4 | 60 | 15.6 | 3.15 ± 0.62 |  |
| 5-6 | 8 | 2.1 | 2.80 ± 0.36 |  |
| 7-8 | 1 | 0.3 | 2.90 ± 0.00 |  |
| 11-12 | 1 | 0.3 | 3.10 ± 0.00 |  |
| Onset of Labour |  |  |  |  |
| Spontaneous | 307 | 79.9 | 2.98 ± 0.64 | t= -0.84; p= 0.40;<br>95% CI -0.22-0.09 |
| Induced | 77 | 20.1 | 3.05 ± 0.52 |  |
| Duration of labour in hours |  |  |  |  |
| 0 – 4 | 59 | 15.4 | 3.17 ± 0.61 | F= 3.69;<br>p= 0.006 |
| 5 – 9 | 143 | 37.2 | 2.97 ± 0.54 |  |
| 10 – 14 | 87 | 22.7 | 2.98 ± 0.55 |  |
| 15 – 19 | 63 | 16.4 | 2.80 ± 0.56 |  |
| 20 – 24 | 32 | 8.3 | 3.19 ± 1.02 |  |
| Mode of Delivery |  |  |  |  |
| Spontaneous vertex delivery | 282 | 73.4 | 2.91± 0.53 | t=-4.84;p<0.001<br>CI=-0.47- -0.20 |
| Caesarean section | 102 | 26.6 | 3.24 ±0.76 |  |

The mean AF size of neonates among groups of duration of labour showed significant difference (*p*<0.05). In the same manner, comparison of AF size in the spontaneous vertex delivery with ceserian section group showed that AF size in the ceserian section group was significantly (p<0.001) higher than spontaneous vertex delivery (Table 4).

The mean AF size of neonates born from mothers of different maternal age (p=0.31), place of residence (*p*=0.31), marital status (*p*=0.91), educational status (*p*=0.18) and monthly income (*p*=0.09) showed no significant difference. However, comparison of the mean AF size of neonates born from mothers of different group in occupation showed statistically significant difference (*p*=0.01) (Table 1).

The size of AF had significant positive correlation (p<0.05) with birth weight and head circumference of neonate (Table 5).

**Table 5.** Correlation of AF size with mean birth weight and head circumference among term neonates of UoGH, Gondar town, Northwest Ethiopia, 2018

| **Correlation of Anterior**<br>**fontanel size (cm)** | **Weight (gm)** | **Head circumference (cm)** |
| --- | --- | --- |
| **N** | 384 | 384 |
| **Mean value** | 3063.50 | 35.70 |
| **Pearson correlation** | 0.11* | 0.17** |
| **Sig.** | 0.04 | 0.001 |

\*. Correlation is significant at the 0.05 level (2-tailed).
\*\*. Correlation is significant at the 0.01 level (2-tailed).
N: 384 in all parameters

## Discussion

Our study focused on the hypothesis that various variables such as sociodemographic factors (maternal age, residency), duration of labour, mode of delivery and the likes contributes for the variability of AF size. Thus, face to face interview using checklist with the mother followed by AF size of apparently healthy term neonates on the first day of life measurement by Popich and Smith method was conducted. Accordingly, gender of the neonate, duration of labour and mode of delivery significantly affects the size of AF. Besides, the birth weight and head circumference of the neonate positively correlated with the size of AF.

In the present study, the mean size of AF of the general population of the study participant is 3.00 ± 0.62 cm. Studies done in Southeastern Nigeria Igbo (16) and black American (17) and calculating the AF size using Popich and Smith method reported similar finding. Interestingly, reports from Nigerian (18), East Indian (19), Hilly Indian (20), Arab Libya (21), white American (17), Chinese (22), Nigerian (23) and Addis Ababa studies (3) were also strongly support our finding. In contrast, it is found to be larger than those reported in Iranian (6), recent Iranian (8), Caucasian American (12), Sri Lankan (24), Israeli (25), Egypt (26), Germany (27), Switzerland (28), Hispanic (29) and Scottish neonates (30). The mean AF size of our study is smaller than in non-hilly Indian (20), Nigerian Ibadan (31) and Indian Nagpur studies (5) (Table 6). The difference in AF sizes could be due to genetic, race or regional (geographic) variation as evidenced from other studies (5, 16, 19, 20). It could also be due to measuring instrument and time of measurement and estimation of AF size in the presence of wide sutures. Wide sutures have been associated with large AF size in the presence of normal head circumference as evidenced by previous studies (16, 32). In some neonates, margins of fontanelle do not meet, but simply merge with suture lines. If tips of a caliper were placed simply at the edges of fontanelle in such neonates, it will result erroneously higher measurement as indicated from previous studies (24, 29). In addition, method difference was noted in Hispanic (presence of premature neonates) and Davies et al. study (got AF size by calculating the surface area) (Table 6). The study in Switzerland (28), Egypt (26) and Germany (27) considered different method to get AF size (Table 6). The method they used makes AF value smaller than from the value in traditional method as explained by previous study (3).

**Table 6.**
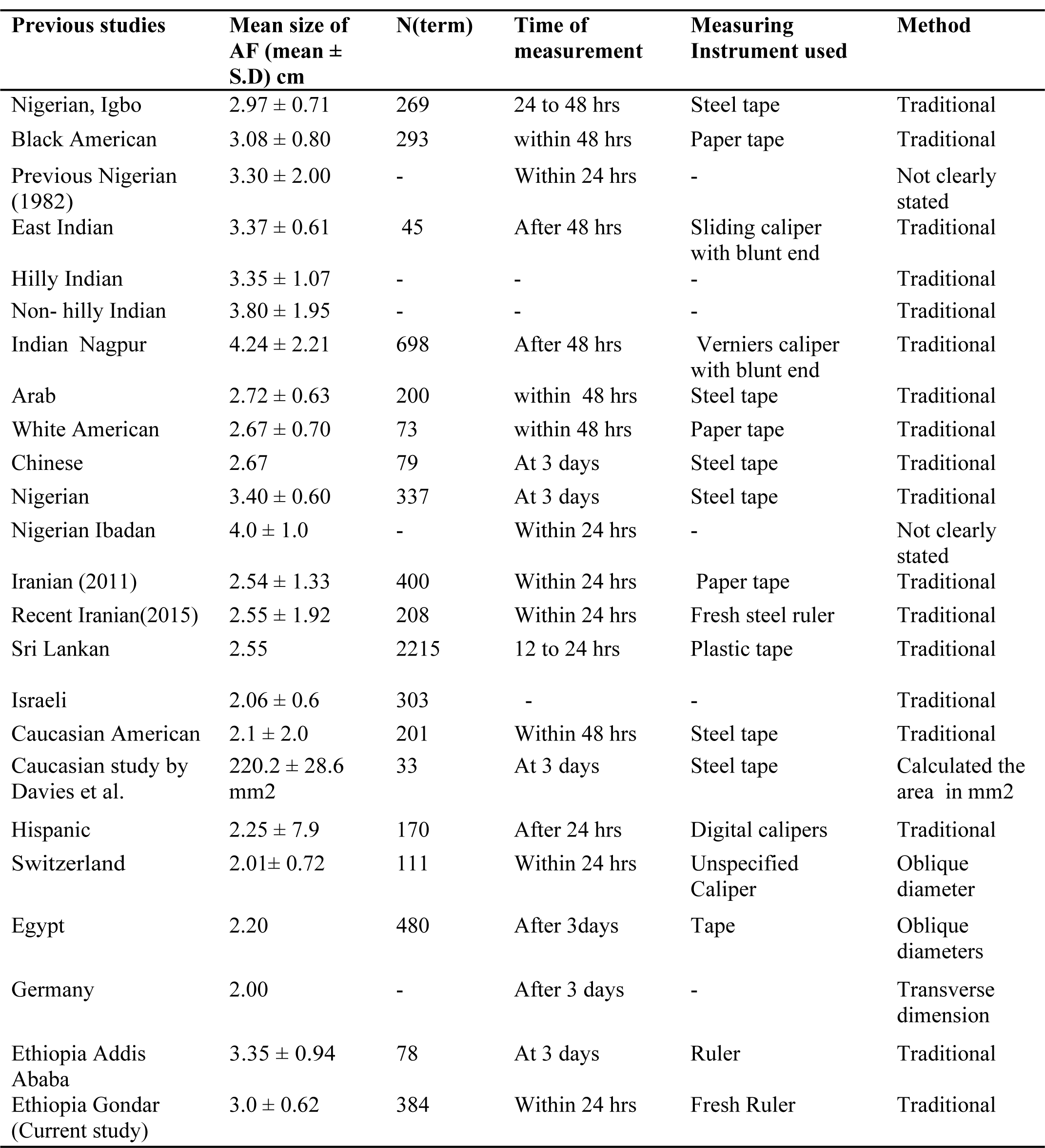
Comparison of mean AF size in neonates of various previous studies

The mean AF size of the present study is comparable with the report from Indian study neonates living in hilly areas; however, it is lower by 0.8 cm from Indian neonates living in non-hilly areas (20). The larger difference in mean AF size indicated that altitude may have an effect on the size of AF (3, 20) but it needs further investigation.

The range of the AF of our study participant neonates is between 1.70 cm and 5.50cm. A study done in Addis Ababa reported AF size was ranged between 1.50cm and 5.80cm. The reason for this slight difference may be related to differences in methodology (timing of measurement and presence of premature neonates).

Piece of evidences reported a variation in AF size between neonates of different races (12, 17, 20, 29), regions (5, 19, 20, 22, 28, 29), genetics (5, 16, 19, 20), nutrition (19), methods, time of measurement, and measuring instruments (16, 29) and estimation in the presence of wide sutures (32). The method, type of instrument measuring and timing differences of various studies are presented in Table 6.

In the present study, the mean size of the male (3.10 ± 0.66 cm) neonates was significantly (*p*≤ 0.05) larger than the female (2.88 ± 0.57 cm) neonates. Our finding is strongly supported by the study conducted among Iranian (6), Arab Libya (21), recent Iranian (2015) (8) and Hispanic (29) study neonates. However, reports from Black and White American (17), Indian Nagpur (5), Scottish (30), Chinese (22), Nigerian Igbo (16), both Caucasian (12, 15), Israeli (25), Nigerian Ibadan (23), Germany (27), Switzerland (28) and Egypt (26) neonates showed no significant difference in AF size between male and female neonates. Regardless of statistically non-significant difference, study conducted in Sri Lankan and Addis Ababa reported that the mean AF size of male neonates is larger than females (3, 24). The discrepancies between our findings and the aforementioned reported studies may be due to differences sample size and timing of measurements as evidenced by previous Hispanic study (29).

In the present study, the mean AF size in different birth order showed barely significant difference (p>0.05). This finding is supported by the study conducted in Addis Ababa (3).

The mean size of AF in different duration of labour showed significant difference (*p*<0.05). There was tendency to decrease in AF size with increasing duration of labour. This finding is supported by the study conducted in Hispanic neonates. The possibility of increased molding secondary to longer labour may play a role as indicated by Hispanic study (29). The mean size of the cesarean section group was significantly (*p*≤0.05) larger than the spontaneous vertex delivery group and the difference may be due to the effect of molding. Our finding is strongly supported by the study conducted in Iran (8). However, reports from previous Iranian (6) and Nigeria Igbo studies (16) showed hardly significant difference in AF size between ceserian section and spontaneous vertex delivery group.

In Ethiopia, different socio-economic status of the mother were compared for AF size in Addis Ababa study (3). Therefore, the present study analyzed for different groups of maternal age, place of residence, marital status, educational status, occupation and monthly income of the mother. All of these variables showed hardly a significant difference (*p*>0.05). But, the materal occupation of the neonate showed significant difference in AF size (p<0.05). The difference may be due to difference level of knowledge to accomplish their daily living activities.

In the present study, a significant positive correlation was found between AF size and birth weight (p<0.05). Our finding is strongly supported by the study conducted among Brazil, Hispanic and Switzerland neonates (28, 29, 33). However, reports from Iranian (6) and Indian Nagpur (5) showed a significant negative correlation between AF size and birth weight. Sri Lankan and Addis Ababa studies reported hardly significant positive correlation between AF size and birth weight (3, 24). Studies conducted among Nigeria Ibadan (31), Nigeria Igbo (16), Scottish (30), and Turkey neonates (14), however, reported that AF size did not have any relation with birth weight.

Significant positive correlation was found between the size of AF and head circumference (p<0.05). Our finding is strongly supported by the study conducted among Brazil (33), Nigeria Ibadan (31) and Addis Ababa (3) neonates regardless of their statistical insignificance. Non-significant correlation (*p*>0.05) between the size of AF and head circumference was reported by the study conducted among Iranian (6), Hispanic (29), Arab (21), Turkey (14), Israeli (25) and Switzerland (28) neonates.

## Conclusion

### The findings of the present study can be summarized as

The mean size of AF for appropriately grown term neonates of the present study had significant difference in mean AF size between males and females. In addition, there was significant difference in mean AF size in neonates of different duration of labour, mode of delivery and occupation of the mother. There was no significant difference in mean AF size in neonates born from mothers of different maternal age, place of residence, marital status, educational status and monthly income of the mother. Furthermore, non-significant difference in neonates of different birth order and onset of labour in mean AF size were found. Besides, the AF size had significant positive correlation with the birth weight and head circumference of neonates. Our finding may, therefore, serve as reference values for pediatricians and other clinicians.

## Acknowledgments

We are grateful to thank the study participant neonates and their mother for their valuable contribution. Our appreciation also goes to Mr. Haimanot G/Hiwot (Assistant Professor of Public Health) for his valuable comments on the statistical analysis. The authors like to express their gratitude to all the members of the Department of Human Anatomy as well as Maternity and Neonatal Ward of the University of Gondar Hospital as their contributions were vital in the completion of this research work. MO acknowledges the receipt of a funding by the University of Gondar.

## Author Contributions

Conceptualization: MO, AT

Formal analysis: MO

Investigation: MO

Methodology: MO, AT

Supervision: AM, AT, EG

Visualization: MO, EG, AT, AM

Writing – original draft: MO, AM, EG, AT

Writing – review & editing: MO, AM, AT, EG

## Supporting Information

**S 1.** Confidentiality and informed consent statement.

**S 2.** Consent form and ethical clearance.

**S 3.** Data collection checklist.

**S 4.** Amaharic version checklist.

